# RNA-ligand interaction scoring via data perturbation and augmentation modeling

**DOI:** 10.1101/2024.06.26.600802

**Authors:** Hongli Ma, Letian Gao, Yunfan Jin, Yilan Bai, Xiaofan Liu, Pengfei Bao, Ke Liu, Zhenjiang Zech Xu, Zhi John Lu

## Abstract

RNA-targeting drug discovery is undergoing an unprecedented revolution. Despite recent advances in this field, developing data-driven deep learning models remains challenging due to the limited availability of validated RNA-small molecule interactions and the scarcity of known RNA structures. In this context, we introduce RNAsmol, a novel sequence-based deep learning framework that incorporates data perturbation with augmentation, graph-based molecular feature representation and attention-based feature fusion modules to predict RNA-small molecule interactions. RNAsmol employs perturbation strategies to balance the bias between true negative and unknown interaction space thereby elucidating the intrinsic binding patterns between RNA and small molecules. The resulting model demonstrates accurate predictions of the binding between RNA and small molecules, outperforming other methods with average improvements of ∼8% (AUROC) in 10-fold cross-validation, ∼16% (AUROC) in cold evaluation (on unseen datasets), and ∼30% (ranking score) in decoy evaluation. Moreover, we use case studies to validate molecular binding hotspots in the prediction of RNAsmol, proving the model’s interpretability. In particular, we demonstrate that RNAsmol, without requiring structural input, can generate reliable predictions and be adapted to many RNA-targeting drug design scenarios.

## Introduction

Drug discovery, a time-consuming and costly process, involves identifying disease-relevant targets and selecting optimal molecules from the expansive chemical space of around 10^60^ drug-like molecules[1, 2]. Currently, most clinical drugs target proteins, yet numerous protein targets are considered “undruggable” due to the lack of suitable structural binding pockets, limiting the range of druggable targets[3, 4]. According to the latest statistics from the DrugBank database[5], merely 854 human proteins have been targeted by FDA-approved drugs. Considering that around 70% of the human genome has the potential to transcribe into RNAs, many of these RNAs exhibit close association with human pathologies, targeting RNA may significantly expand the pool of druggable targets. Originating with ribosomes as crucial antibiotic targets[6-8], RNA-targeting has burgeoned in the last decade, various RNA types including mRNA, miRNA, tRNA, rRNA, and long non-coding RNAs (lncRNAs) have been proved to be targets of small molecules [9-15]. Most of the well-known RNA-targeted small molecules are identified using phenotypic screening occasionally, for instance, Evrysdi (risdiplam)[9, 16], approved by the FDA in August 2020, targets human mRNA, correcting specific splicing defects in treating spinal muscular atrophy. Moreover, ribocil, a small molecule targeting FMN riboswitches is pivotal in bacterial regulation and antibiotic resistance[10]. These experiences suggest the transformative potential of RNA-targeting in the field of drug discovery. Currently, researchers have applied target-based high-throughput screening (HTS) techniques derived from protein-targeting drug discovery[17-19] such as the automated ligand identification system (ALIS) and small-molecule microarrays (SMM) to identify RNA-binding small molecules[20, 21]. For example, using ALIS, the compound X1 was identified to bind to the lncRNA Xist, inhibiting X chromosome inactivation by inducing conformational changes that disrupt its interaction with associated protein factors[11]. Also, a recent work used SMM to screen large libraries of compounds against a set of disease-related RNA targets and collected the largest fully public nucleic acid binding small molecule library named Repository Of Binders to Nucleic acids (ROBIN)[22].

However, since existing experimental methods are costly and labor-intensive, many computational methods have been proposed as alternative solutions to automate the identification of RNA-targeting small molecules. Firstly, many methods collected existing experimental validated RNA targets and RNA-binders into libraries and predicted RNA-small molecule binding by assessing the similarity between query data and curated data in library, such as Inforna [20, 23], RNAligands [24], and RSAPred [25]. Secondly, for RNA targets of interest with known structures, molecular docking remains the most straightforward virtual screening method, several docking and scoring methods have been developed for RNA-targeting ligands, such as rDock[26], RLDOCK[27], AutoDock Vina[28]. Despite the widespread use of molecular docking, its accuracy is limited due to factors such as force field settings, inaccuracies in scoring functions[29, 30], and inadequate sampling of ligand conformations[31]. Thirdly, many studies have begun to utilize advanced deep learning models to study RNA-ligand interactions. These studies roughly fall into three categories: predicting small molecule binding sites on RNA target structures (site model)[32-35], designing potential binding ligands for RNA structural pockets (generative model)[36-38], and predicting RNA-ligand binding interactions (classification model)[39]. Site models were proposed to predict the positions or local regions on the RNA target as binding sites/motifs by the representation and characterization of multiple properties for 3D structures of RNA targets. Generative models began with the RNA pocket, using deep learning models to design the candidate ligand for given RNA pockets. For example, RNAmigos and RNAmigos2 models use the augmented base pairing network (ABPN) representation of 3D RNA pocket structure and use a relational graph convolutional neural network module to generate the fingerprint of potential binding ligand. Classification models were developed to leverage the combination of RNAs and ligand features for predicting RNA-ligand interactions.

Despite all these efforts, aforementioned library-based methods depend on in-house experimental databases and exhibit poor generalizability on unseen queries. Current computational models heavily rely on RNA 3D structure information, while there are only 7,806 RNA-containing structures in the RCSB Protein Data Bank (PDB)[40] (http://www.rcsb.org/), accounting for around 3.5% of the total number of structures (221,371 as of Jun 2024). Moreover, many disease-related human mRNAs[12, 41] and lncRNA targets (e.g., XIST [11], MALAT1[42], HOTAIR[43]) lack defined structures or have structures that are difficult to determine[44-46], making them unsuitable for the aforementioned methods as input or for training. Given the widespread application of deep learning technology in predicting protein sequence-ligand binding[47-55] and RNA sequence-protein binding[56, 57], it is feasible to leverage state-of-the-art deep learning models to establish a sequence-based RNA-small molecule prediction method for RNAs with unknown structures. Besides, recent structure-based virtual screening (SBVS) methods for protein targets[58, 59] have attempted to improve prediction performance on unseen data. Currently, no RNA-targeting models have systematically proven their ability to generalize on unseen datasets. Although the existing methods demonstrate promising performance in traditional model evaluations, determining the binding pattern of RNA and small molecules while simultaneously accelerating the development process of RNA-targeting small molecule drugs remains beyond our current capabilities. Furthermore, there is ample room for improvement in the interpretation and adaptability of existing models.

To address these challenges, we present RNAsmol, a novel sequence-based RNA-small molecule interaction scoring model for RNAs with unknown structures. We integrated diverse information from heterogeneous data sources including PDB and ROBIN and carefully preprocess these datasets to disclose and interpret the binding between RNA and small molecules. Leveraging data perturbation and augmentation strategies, RNAsmol aims to address bottlenecks such as data scarcity, comprehensively characterize the binding patterns between RNA and small molecules thereby aiding the development process of small molecule drugs targeting RNA. We utilized graph diffusion convolution for molecular feature representation and bilinear attention feature fusion modules to predict RNA-small molecule interactions. Then, we employed four evaluation methods with progressively stricter assessments to benchmark RNAsmol. RNAsmol achieved significant performance compared to other methods, showing an average improvement of approximately 8% in ROCAUC during 10-fold cross-validation, around 16% in ROCAUC for cold evaluations on unseen datasets, and about 30% in ranking score during decoy evaluations. Furthermore, we validate the model’s interpretability through case study validations, identifying molecular binding hotspots corresponding to RNAsmol’s predictions. For structured molecules like most proteins and certain noncoding RNAs (e.g., Riboswitch and Ribozyme), there are many AI-driven methods available. However, for RNAs without stable tertiary structures (e.g., many mRNAs and lncRNAs), there is still a lack of prediction methods for RNA-ligand interaction scoring. RNAsmol is capable of generating reliable predictions without relying on structural input, can be applicable to various RNA-targeting drug design scenarios.

## Results

### Overview of RNAsmol framework

As illustrated in **Figure 1**, we build a deep learning model termed RNAsmol which takes RNA sequences and small molecules as inputs and outputs the likelihood of their binding as binding score. To address the issues arising from data scarcity and learning biases, as shown in **Figure 1a**, we apply three kinds of data perturbations on the raw RNA-small molecule interaction network in our study, i.e., *ρ*_*r*_ for RNA perturbation, *ρ*_*m*_ for small molecule perturbation, *ρ*_*n*_ for interaction network perturbation. RNA perturbation adds the shuffled RNA sequences with same dinucleotide frequency as the RNA targets into the raw network, small molecule perturbation adds drug-like compounds with high MACCS fingerprint similarity to the small molecule ligands into the raw network, and the interaction network perturbation introduces negative labels in the unknown interaction space of the raw network. Along with three kinds of augmentation strategies for each perturbation, we generate three kinds of training datasets, i.e., *T*_*r*_, *T*_*m*_, *T*_*n*_ for the model. **Figure 1b** shows the overall model architecture of RNAsmol, we utilize parallel processing modules for RNA targets and their corresponding small molecule ligands. Specifically, we employ a multi-view convolutional neural network for RNAs which strengthens long-range context aggregation for comprehensive representations, and a graph diffusion convolutional neural network for small molecules which extracts global topological properties of molecular structures. Then we utilize a bilinear attention block as a feature fusion module further aiding in annotating key binding sites relevant to their interactions and a multilayer perceptron (MLP) for classification in the model. To prove the prediction performance and robust model generalization and interpretability, as shown in **Figure 1c**, we sequentially use the 10-fold cross-validation (CV) evaluation, cold evaluation, decoy evaluation and case study validations to compare the performance of RNAsmol with other models, with each subsequent evaluation introducing progressively stricter criteria. Besides, we also refine the parameters in RNAsmol in post-hoc analysis and optimize the model as a tool for drug virtual screening. See **Methods** for more details about RNAsmol model modules and evaluation methods.

**Figure 1.**
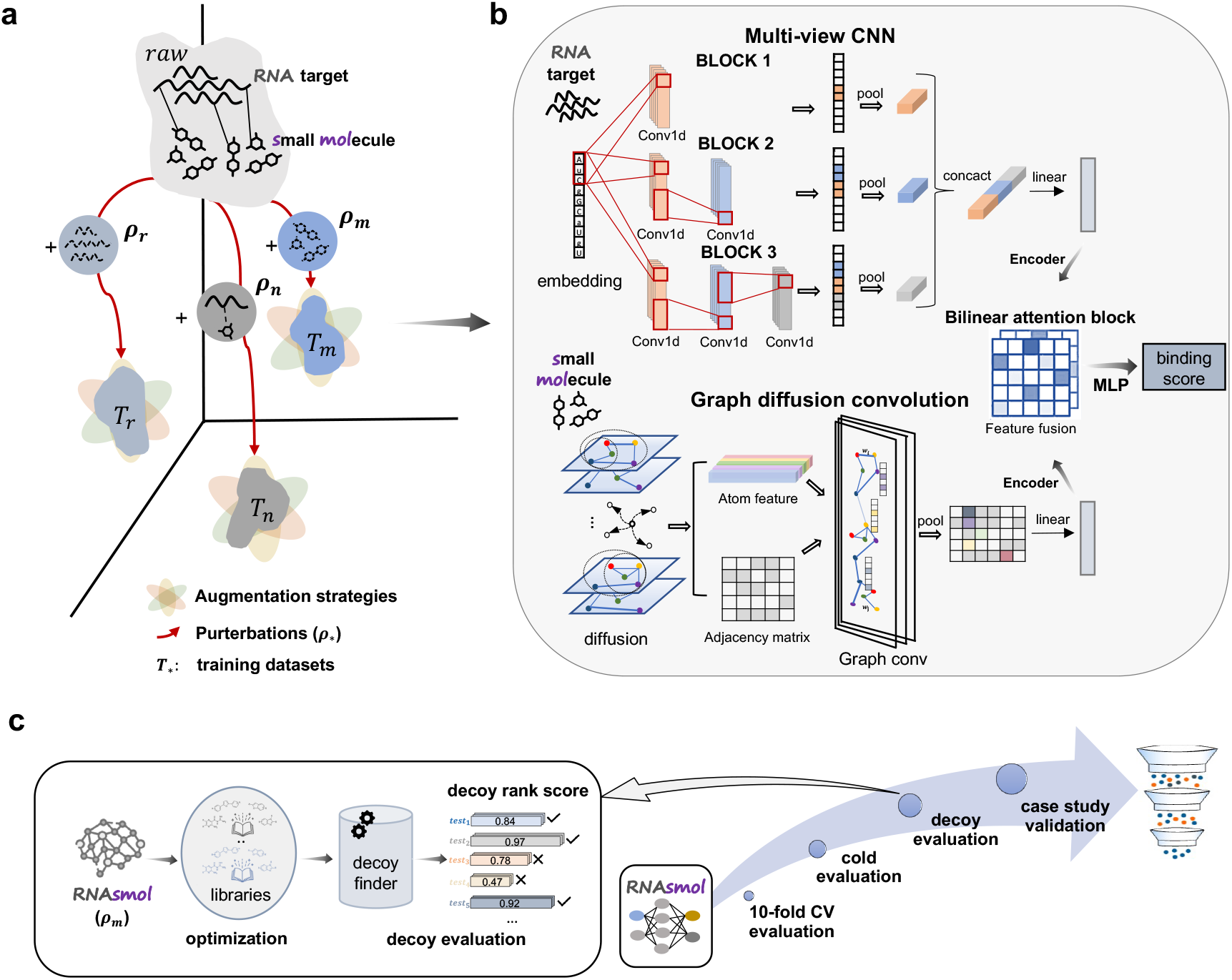
Overview of RNAsmol framework. **a**. Three kinds of perturbations with augmentations on RNA-small molecule interaction network. **b**. Model architecture. RNAsmol model has two parallel feature extraction modules, multi-view CNN for RNA target and graph diffusion convolution for small molecule respectively. Then, it employs a bilinear attention block for feature fusion and a multilayer perceptron (MLP) for classification. **c**. Evaluations and reliable post-hoc analysis of RNAsmol model. Four kinds of evaluations including 10-fold CV (cross-validation) evaluation, cold evaluation, decoy evaluation and case study validation are utilized to prove the reliable performance on classification task and robust potential on drug virtual screening. Additionally, we optimize small molecule perturbation for decoy evaluation.

### RNAsmol provides accurate predictions of RNA-small molecule binding in perturbation space

As a binary classification model for predicting RNA-small molecule interactions, we compared RNAsmol with four recent sequence-based target-drug interaction prediction models: MGraphDTA_RNA, IIFDTI_RNA, GraphDTA_RNA and DrugBAN_RNA (see **Methods** for details). Firstly, we evaluated the prediction performance of RNAsmol with three types of augmentations against these models in perturbation space, as shown in **Figure 2a**. The x, y, and z axes represent three types of perturbations, and each point’s coordinates in this 3D scatter plot correspond to the average ROCAUC or PRAUC values of a model based on a 10-fold CV under a specific perturbation. The confidence ellipses for the five models suggest that RNAsmol robustly outperforms the other models across all perturbation settings on both the PDB and ROBIN datasets. Besides, *ρ*_*m*_ enhance the best predictions for RNAsmol and MGraphDTA_RNA, while other models fail to achieve a steady prediction state within the perturbation space. Compared to the ROBIN dataset, the performance on PDB dataset has higher variations within the perturbation space. Secondly, to disclose the effectiveness of data augmentation strategies, we compare eight models, including MGraphDTA_RNA, IIFDTI_RNA, GraphDTA_RNA and DrugBAN_RNA, as well as RNAsmol with and without data augmentations (RNAsmol_noaug, RNAsmol_rnaaug, RNAsmol_molaug and RNAsmol_bothaug) (see **Methods** for details) on three kinds of perturbations across all metrics of 10-fold CV evaluation, including ROCAUC, PRAUC, ACC, SEN, SPE, and F1 score, as illustrated in **Figure 2b**. Using the Mann-Whitney-Wilcoxon test with Bonferroni correction, the p-values indicate that all RNAsmol models outperform the other models, and data augmentation strategy significantly improves the predictive performance of our model. Since augmenting both RNAs and small molecules achieves the best prediction across all perturbations, we selected RNAsmol_bothaug for subsequent evaluations and comparisons. Thirdly, to evaluate and compare RNAsmol with other models of ROCAUC with 10-fold CV and cold evaluations in which **we conducted cold evaluation** for RNA targets, small molecules, and both interaction molecules (see **Methods** for details). As shown in **Figure 2c** and **Figure S1**, our model consistently outperformed the other models in four kinds of settings, demonstrating superior robustness in the context of unseen evaluations. RNAsmol outperforms other methods with average improvements in ROCAUC of 0.12 on the PDB dataset and 0.05 on the ROBIN dataset in 10-fold cross-validation. In cold evaluation settings, it shows improvements of 0.2 on PDB and 0.11 on ROBIN for cold evaluation on RNA, 0.16 on PDB and 0.07 on ROBIN for cold evaluation on small molecules, and 0.3 on PDB and 0.15 on ROBIN for cold evaluation on RNA-small molecule pairs. The results indicate that when both interacting molecules were unseen during training, the model’s predictions were most affected, followed by unseen RNA molecules, with the least impact observed when small molecules were unseen. Although all models show variable predictions on the PDB dataset from **Figure 2a**, our model demonstrates a higher improvement than other models on the PDB dataset than ROBIN dataset in both 10-fold CV and cold evaluations.

**Figure 2.**
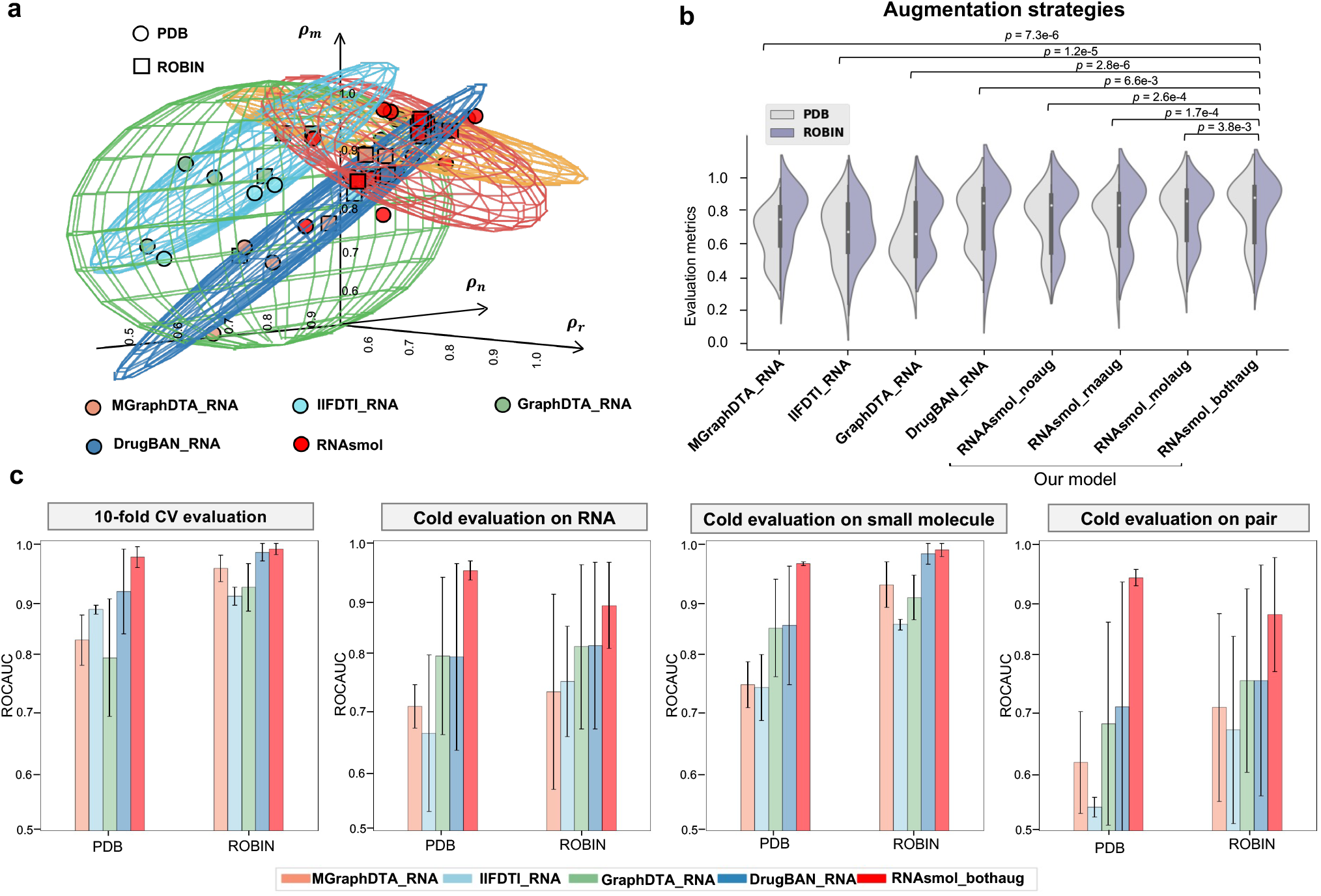
Performance comparison in predicting RNA-ligand interaction based on 10-fold cross-validation (CV) and cold evaluation. **a**. Predictions of five classification models across two RNA-small molecule binding datasets on perturbation space. **b**. Comparisons between RNAsmol with three kinds of augmentations and other models of evaluation metrics including ROCAUC, PRAUC, ACC, SEN, SPE, F1 score, p-values are obtained from the Mann-Whitney-Wilcoxon test with Bonferroni correction. **c**. Comparison with other models on 10-fold CV evaluation and three kinds of cold evaluation strategies (test on unseen data). Error bar represents the standard deviation (STD) calculated from multiple folds and perturbations.

### RNAsmol provides reliable and adaptable predictions with molecular perturbation (*ρ*_*r*_ and *ρ*_*m*_)

To demonstrate the extensive application and suitable scenarios of RNAsmol with RNA perturbation and small molecule perturbation, we conduct cross-RNA type test on PDB dataset and target-specific predictions on ROBIN dataset. The PDB dataset encompasses various RNA types, including rRNA, riboswitch, viral RNA, ribozyme, aptamer, primer complex, and splicing-related RNAs. We found that the interaction networks of PDB and ROBIN datasets exhibit different properties. Furthermore, as illustrated in **Figure 3a**, the calibration curves indicate that the model’s predicted binding scores are consistent with actual outcomes, demonstrating that RNAsmol (*ρ*_*r*_) is well-calibrated on the PDB dataset. As shown in **Figure S2**, cross-dataset tests (where the training set is PDB and the test set is ROBIN, or *vice versa*) revealed that these two datasets cannot predict each other effectively. This performance decrease from within-dataset tests (where both training and test sets are either PDB or ROBIN) is more pronounced under RNA perturbation (*ρ*_*r*_) conditions, indicating significant differences in RNA target profiles between the datasets. Interestingly, due to the substantial overlap in the physicochemical properties of small molecules in both PDB and ROBIN datasets (**Figure S17**), small molecule perturbation models are more robust in cross-dataset predictions, resulting in less performance decline. This suggests that small molecule perturbation models maintain their predictive performance across different datasets, whereas RNA perturbation models face greater challenges. However, the pronounced decrease in performance of RNA perturbation models in cross-dataset tests indicates their sensitivity to capturing binding signals within RNA-small molecule interaction networks with different RNA profiles. To leverage this sensitivity, we apply the RNAsmol (*ρ*_*r*_) to explore predictions across different RNA types and their cross-dataset predictions. Our cross-RNA type test results, shown in **Figure 3b**, reveal that RNAsmol (*ρ*_*r*_) performs best on riboswitch targets in within dataset prediction and also generalizes well to other RNA types in cross-dataset prediction. Therefore, the RNA-specific features captured by the RNAsmol generalize well on dataset with a shift in distribution of RNAs’ properties. Besides, these findings suggest that RNA perturbation models are particularly effective in capturing the nuanced interactions within the PDB dataset, making them a valuable tool for RNA type-specific drug virtual screening There are 27 disease-related RNA targets in ROBIN dataset, according to the screening results, as illustrated in **Figure 3d**, the 27 RNA targets are grouped into five kinds of secondary structure: RNA G-quadruplex (rG4), hairpin, pseudoknot, three-way junction and triple helix. Structures such as rG4s, pseudoknots and three-way junctions exhibit the highest selective hit rates which refers to the proportion of small molecules exclusively hitting a target without hitting others, as indicated by the stacked bars. To investigate RNA-small molecule interactions for individual RNA target with high selective hit rate and different secondary structure in the ROBIN dataset, we trained RNAsmol with small molecule perturbation on single RNA target. As shown in **Figure 3e**, RNAsmol (*ρ*_*r*_) performs well across all RNA targets, making it suitable for prefiltering compound libraries before screening experiments. The rG4 targets, including EWSR1, AKTIP, demonstrate higher sensitivity and recall among these targets which means the RNA-binders of these targets can be sensitively and well detected. Meanwhile, pseudoknot targets such as ZTP and three-way junction targets such as TPP and Glutamine_RS exhibit higher specificity and precision, indicating that although the RNA-binders for these targets may not be detected as frequently, the detections are very reliable when they occur. The average dissimilarity among high-specificity predictions is greater than that among high-sensitivity predictions, suggesting the diverse prediction patterns of RNAsmol (*ρ*_*m*_) on individual RNA targets. Additionally, RNAsmol (*ρ*_*m*_) makes calibrated and accurate predictions for both the full ROBIN dataset and individual RNA target in ROBIN dataset, as indicated by the calibration curve aligning well with the actual probabilities (**Figure 3c** and **Figure S5**).

**Figure 3.**
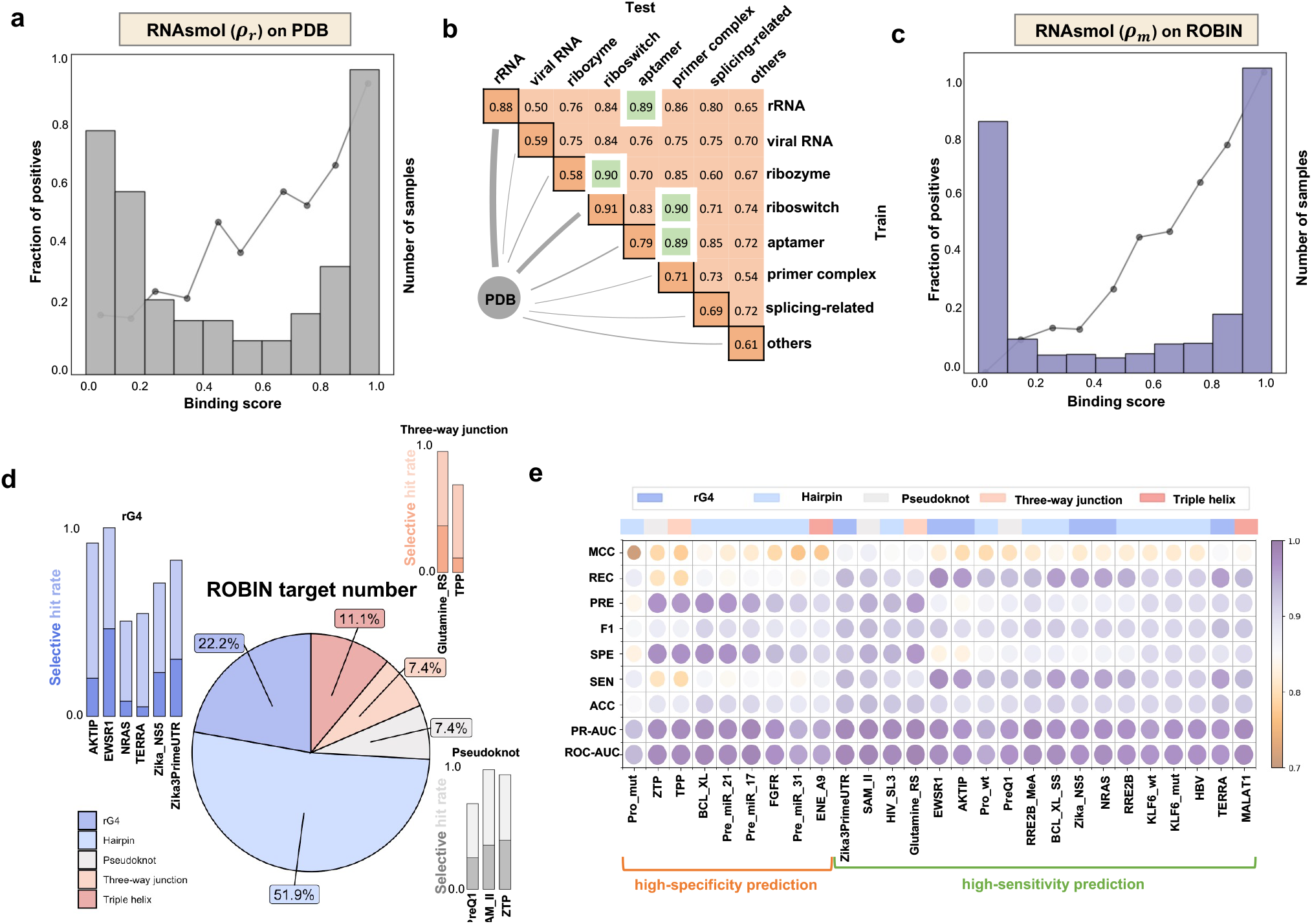
Applications of small molecule perturbation (*ρ*_*m*_) and RNA perturbation (*ρ*_*r*_) on PDB and ROBIN datasets. **a**. Calibration curve of RNAsmol (*ρ*_*r*_) classification on PDB datasets. b. ROCAUC heatmap of the ROCAUC in cross-dataset evaluation within and across RNA types in the PDB dataset. The rows represent the training dataset and the columns represent the test dataset. c. Calibration curve of RNAsmol (*ρ*_*m*_) classification on ROBIN datasets. d. RNA target numbers categorized by different structures in the ROBIN dataset. Stacked bar charts depict the hit rate and selective hit rate of rG4 (RNA G-quadruplex), pseudoknot, and three-way junction targets in screening experiments. Hit rate refers to the proportion of small molecules hitting each target, while selective hit rate indicates the proportion of small molecules exclusively hitting a particular target without hitting others. **e**. Average metrics including ROCAUC, PRAUC, ACC, SEN, SPE, F1, PRE, REC, MCC of 10-fold CV for individual target in the ROBIN dataset. High-specificity and high-sensitivity predictions are stratified according to the hierarchical clustering result.

### Optimization of the small molecule perturbation (*ρ*_*m*_) for decoy evaluation

Given that RNAsmol (*ρ*_*m*_) provides the most robust prediction in classification tasks (see **Results** Section 2 for details), we aim to use the binding score predicted by RNAsmol (*ρ*_*m*_) as a constraint to narrow down the vast drug-like chemical space. There are several drug-like compound libraries used for high-throughput drug screenings, including the ZINC bioactive compound library, COCONUT natural product (organic molecules) library, ChemBridge BuildingBlocks (chbrbb) library and BindingDB protein binder library. **Figure 4a** shows the UMAP visualization of the molecular physicochemical properties including molecular weight (MW), partition coefficient (logP), hydrogen bonds donors (HBD), hydrogen bond acceptors (HBA) and the number of rotatable bonds (RB) across these four drug-like compound libraries, the PDB dataset and the ROBIN dataset. Clustering results reveal that molecules from the BindingDB database exhibit higher similarity to RNA-binder molecules in the two datasets in terms of physicochemical properties. Conversely, molecules in the chbrbb library display the most divergent distribution properties, while those in the COCONUT library demonstrate the most extensive range of molecular physicochemical properties. We then employed 10-fold CV evaluation and decoy evaluation to investigate and optimize the small molecule perturbation using three different background compound libraries. **Figure 4b** shows that RNAsmol (*ρ*_*m*_) get the best classification performance when using the largest COCONUT libray and the worst performance on BindingDB dataset. However, employing the aforementioned three small molecule datasets as background drug libraries, and the bioactive small molecules from ZINC as the decoy drug library, as shown in **Figure 4c**, RNAsmol (*ρ*_*m*_) achieved optimal decoy evaluation results when utilizing molecules from the BindingDB database as the background. Furthermore, small molecule ligands binding to RNA targets tend to exhibit selectivity, and the chemical property space of RNA ligands overlaps to some extent with protein ligands. Therefore, we infer that using a negative dataset composed of molecules with similar physicochemical properties during model training enables the acquisition of more precise features for distinguishing drug-like molecules.

**Figure 4.**
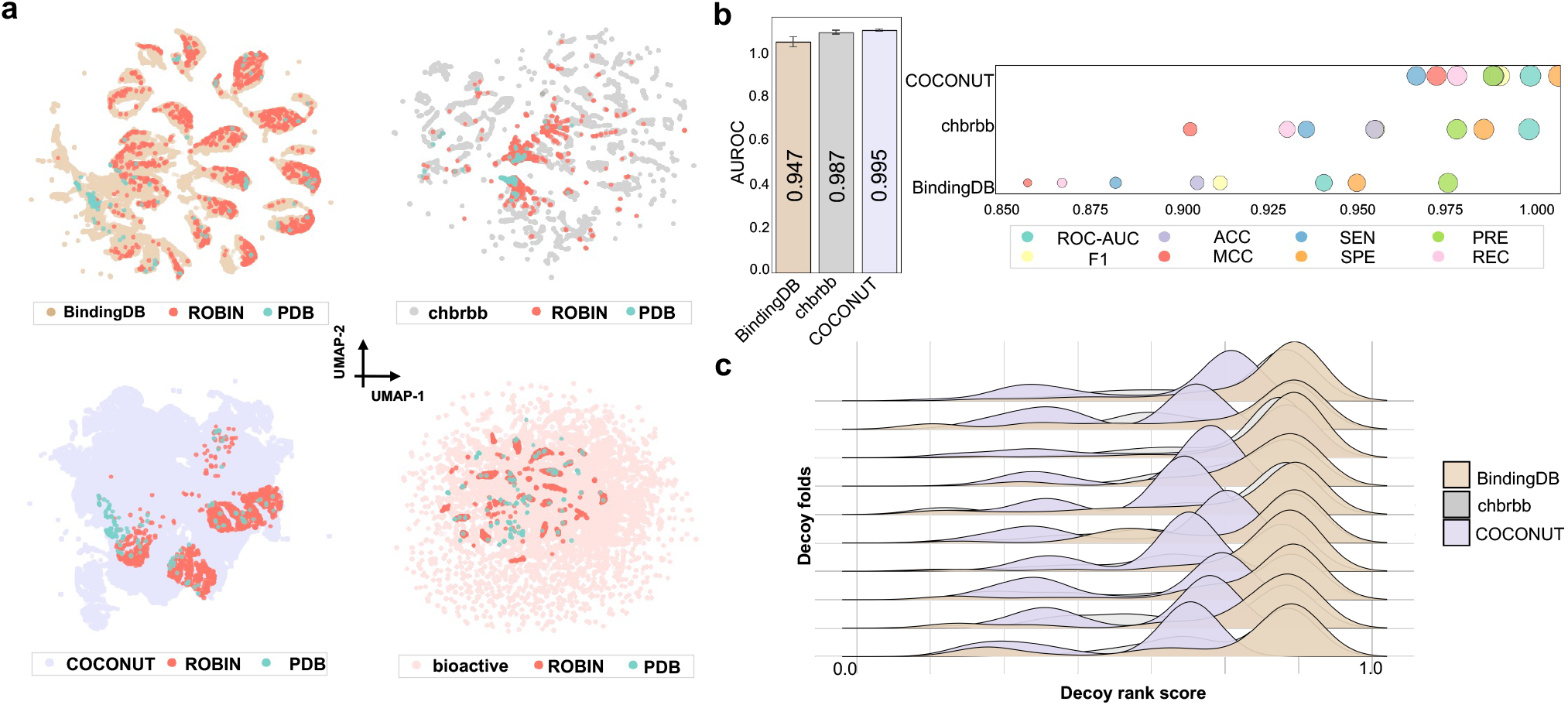
Optimization of the small molecule perturbation (*ρ*_*m*_) for decoy evaluation. **a**. UMAP visualizations of molecular physicochemical properties including molecular weight (MW), partition coefficient (logP), hydrogen bonds donors (HBD), hydrogen bond acceptors (HBA) and the number of rotatable bonds (RB). The first three plots show the overlap between RNA-binding small molecules (from both PDB and ROBIN) and drug-like background compound libraries (BindingDB, chbrbb, and COCONUT), while the fourth plot shows the overlap with bioactive small molecule libraries used in decoy set generation. **b**. Classification comparison of RNAsmol (*ρ*_*m*_) with three drug-like background compound libraries. Error bar in barplot represents the standard deviation (STD) calculated from 10 folds. **c**. Comparison of decoy rank score distribution of RNAsmol (*ρ*_*m*_) with three drug-like background compound libraries. Higher decoy rank score indicate the better performance in decoy evaluation.

### RNAsmol effectively distinguish known RNA-targeting small molecule from decoys

First, we trained the aforementioned model, selecting RNAsmol (*ρ*_*m*_) with the optimized background library, i.e., BindingDB. Subsequently, for each RNA target in the test set of PDB dataset, we generated a decoy evaluation set consisting of bioactive small molecules in ZINC bioactive small molecule library using decoyfinder[60] software. We used the trained RNAsmol model to predict binding scores for each small molecule in the decoy set and get the rank of true ligand in the predicted binding scores of decoy set. Similarly, we employed the RNAmigos model to generate a molecular representation vector, calculated the distance between this vector and the fingerprint of both true and decoy small molecules, and then ranked the results. Then we use RNAmigos2 model and rDock software to get a score for each decoy molecule and get the rank of true ligand in the decoy set. Finally, we compared the ranking outcomes of the four models, as depicted in **Figure 5a**, the boxplot illustrates the distribution of rankings for positive small molecules in the 10-fold decoy test. Notably, our model’s rankings significantly outperformed those of the other three models, achieving an average decoy rank score of 83%, which was 45% higher than RNAmigos, 6% higher than RNAmigos2, and 40% higher than rDock. As shown in **Figure 5b**, upon randomizing RNA targets, our model exhibited greater variation in ranking distribution measured by Kullback-Leibler (KL) divergence, indicating its superior specificity for RNA targets. We calculated fingerprint similarity using four distance metrics: Euclidean distance, cosine distance, Chebyshev distance and correlation distance, with differences shown in **Figure S8**. For RNAmigos2, we used four modes of this model including dock mode, native mode, fp mode and mixed mode for the evaluation, and the corresponding results are also shown in **Figure S8**. Besides, we also apply two trained RNAsmol models which are trained on PDB and ROBIN datasets respectively on many cases as a drug virtual screening application. As shown in **Figure 5c**, RNAsmol_ROBIN have higher performance on new-revealed RNA-targeting drugs like Ribocil, Risdiplam, etc, while RNAsmol_PDB perform better on riboswitch cases. RNAsmol makes prediction on RNA targets which are with unknown structure and has overall better performance than the other RNA-targeting virtual screening models including RNAmigos, RNAmigos2 and RSAPred_riboswitch.

**Figure 5.**
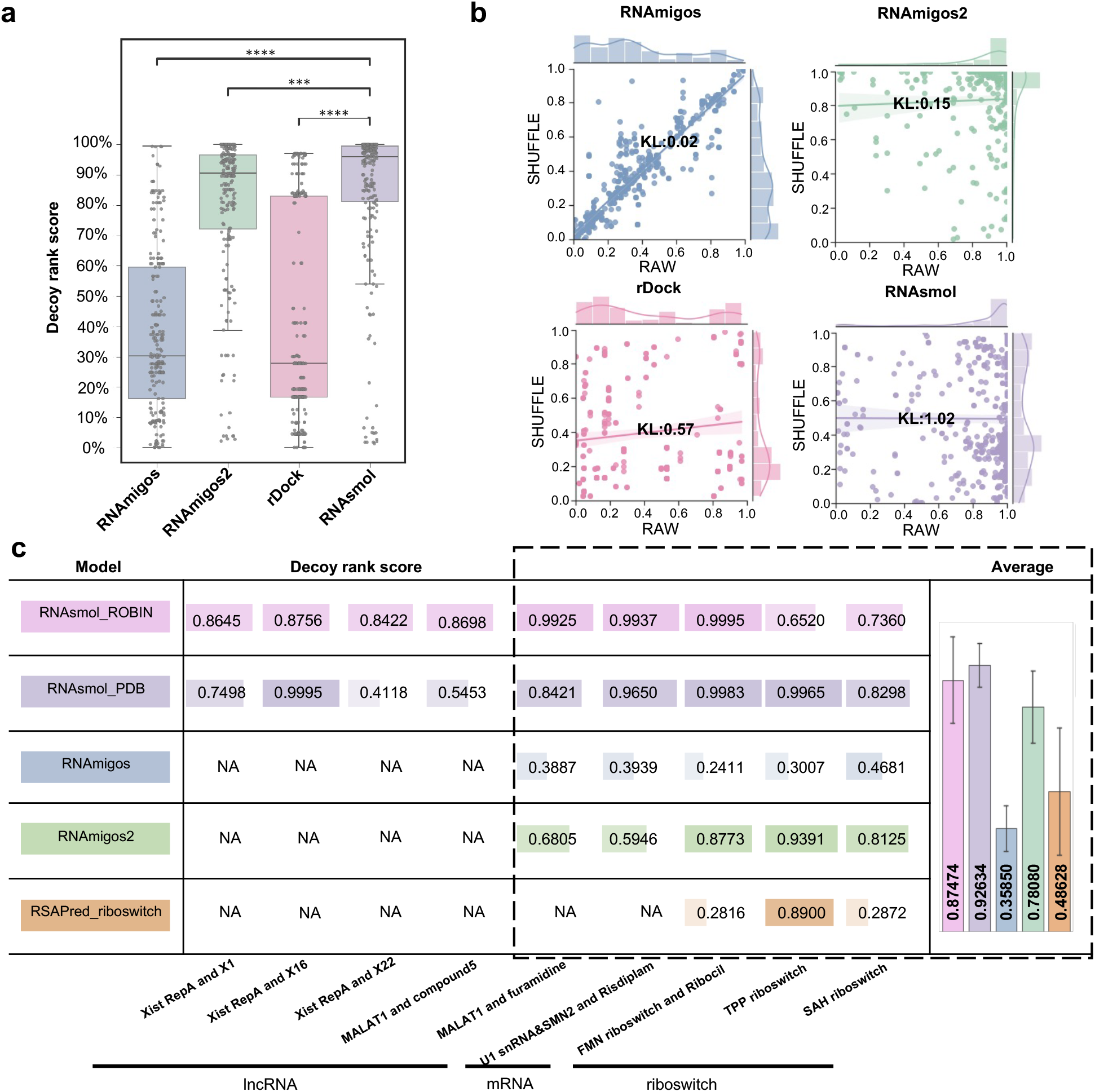
Performance comparison in virtual screening based on the decoy evaluation. **a**. Decoy rank score performance comparisons with other structure-based methods of decoy evaluation on PDB dataset. * P-value<0.05, ** P-value<0.01, *** P-value<0.001, **** P-value<0.0001, Wilcoxon rank sum test, one-tailed (RNAsmol has higher decoy rank score than RNAmigos, RNAmigos2, rDock). **b**. Decoy rank score distribution of RNAsmol and other methods with and without RNA target shuffle. Kullback–Leibler (KL) divergence measures the difference between the decoy rank score distribution of RAW and SHUFFLE (higher KL values indicate greater differences) **c**. Decoy rank score comparisons between two trained RNAsmol models (trained on ROBIN dataset and PDB dataset separately) and other models on well-known RNA-targeting drug cases.

### RNAsmol provides interpretable predictions of RNA-small molecule interaction

To further validate the interpretability of the model, we visualized the hotspots on RNA and small molecules by gradient-weighted class activation mapping (Grad-CAM). **Figure 6** displays the structure of class I pre-queuosine1 (PreQ1) riboswitch from *Bacillus subtilis* (PDB ID: 3K1V) and ZTP riboswitch from *Fusobacterium ulcerans* (PDB ID: 5BTP) respectively. One the left of **Figure 6a** and **Figure 6b**, present the structural snapshots of two riboswitches binding to small molecules, as rendered in PyMOL, the hydrogen bonds between RNA and ligand are colored light blue with the annotated distance, and the middle part shows the profile of contacts generated by Ligplot+ software. Then we employed the Class Activation Map (CAM) module to obtain the weights of the last convolutional layer in the convolutional neural network through backpropagation. Subsequently, these weights were multiplied with the feature map of that layer to obtain a weighted sum, forming a feature map. This enabled the mapping back to atoms in small molecule and RNA target to visualize the importance of each atom or nucleic acid for classification (**Figure S9**). On the right part, we showcased the weights on the small molecule binding to the PreQ1 riboswitch and the pseudoknot motif in ZTP riboswitch. The regions in the model with higher weights often correspond to key atoms or nucleic acid involved in binding in real structure, indicating that our model demonstrates a high consistency in predicting hotspots on small molecules and regions where hydrogen bonds are formed. Results of the visualizations of c-di-GMP-II and S-adenosylhomocysteine (SAH) riboswitches are shown in **Figure S10** and **Figure S11** respectively.

**Figure 6.**
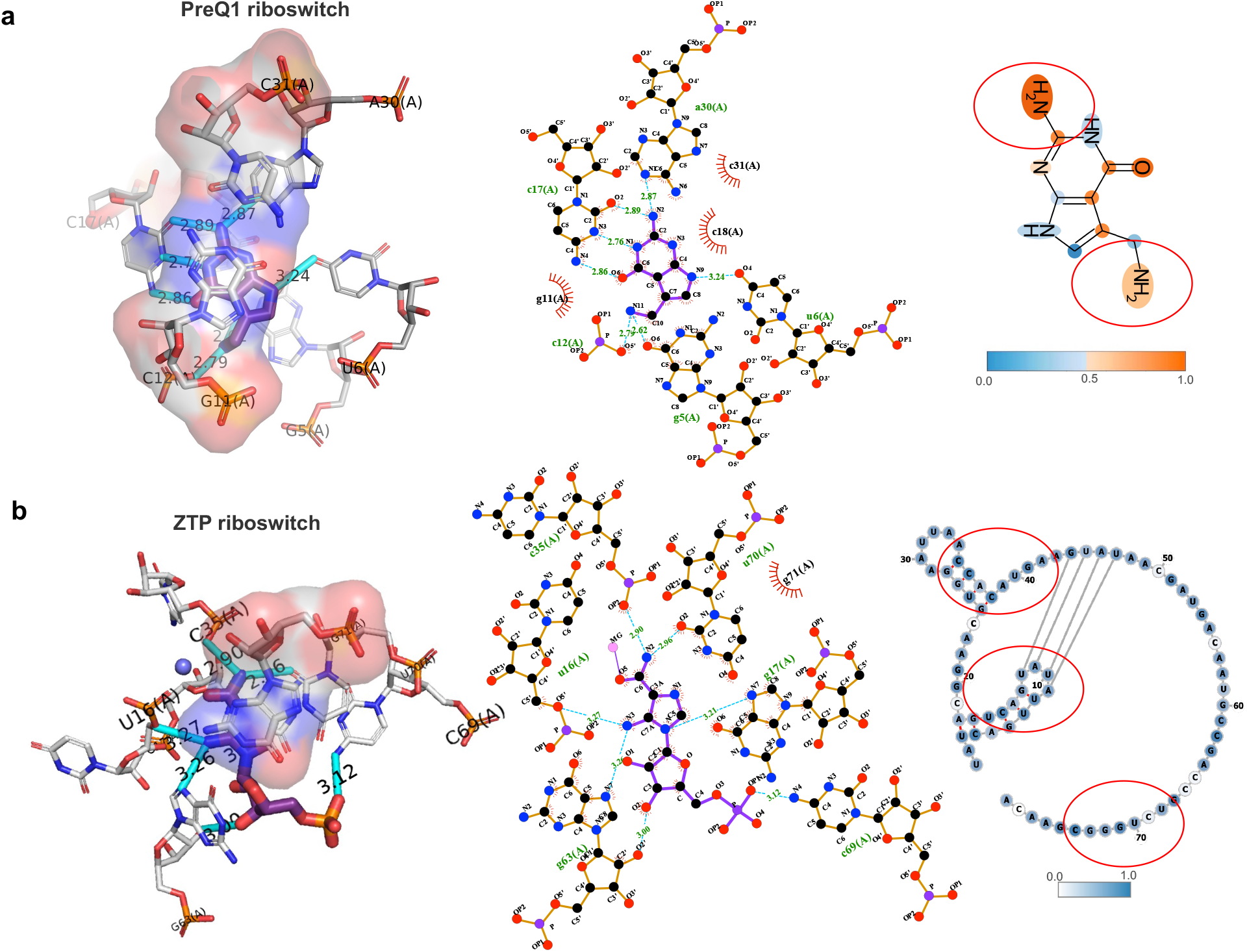
Case study validation and visualizations of molecular hotspots of RNAsmol prediction. a. Structural snapshots of class I pre-queuosine1 (PreQ1) riboswitch from *Bacillus subtilis* (PDB ID: 3K1V) structure by PyMOL, profile of contacts within the binding site by ligplot software, and Grad-CAM weight visualization of PreQ1 ligand in RNAsmol prediction. Hydrogen bonds are colored light blue with annotated distance in both the structure and profile. b. Structural snapshots of ZTP riboswitch from *Fusobacterium* ulcerans (PDB ID: 5BTP) structure by PyMOL, profile of contacts within the binding site by ligplot software, and Grad-CAM weight visualization of ZTP target secondary structure in RNAsmol prediction. Hydrogen bonds are colored light blue with annotated distance in both the structure and profile.

## Discussion

To summary, AI-driven RNA-targeting drug design would provide crucial insights for the development of targeted therapeutics. We proposed a unified framework for RNA-ligand interaction scoring via data perturbation and augmentation modeling. Through comprehensive testing across multiple evaluations, we demonstrate superior performance of our model compared to existing ones. Additionally, we conduct discussions on different applications on various RNA-targeting drug design and drug screen senarios, aiming to elucidate patterns and preferences in the interaction between RNA and small molecules. We proved that the sequence input did not introduce significant noise into our model. Instead, our data perturbation and augmentation strategies successfully enriched the informative content within the sparse data space of RNA-small molecule interactions. This approach significantly enhances our understanding of how RNA interacts with small molecules. Moreover, beyond achieving strong predictive performance in binding prediction, our model excelled particularly in the strictest decoy evaluations. Decoy evaluations challenge the model to distinguish accurately between true and false molecules in unseen but similar datasets using trained RNAsmol model. This success can be attributed to our meticulous approach in selecting and preprocessing the existing RNA sequence containing binding sites from chains and the stringent selection of drug-like small molecules, which closely mirrors real-world drug screening scenarios. By optimizing small molecule perturbations, we gained valuable insights into the nuanced properties of RNA-binders within the drug-like chemical space, thereby contributing to our robust performance in decoy evaluations. In contrast, pocket-guided SBVS models not always exhibit target specificity as evidenced by RNA target shuffle decoy results. Our model has effectively learned the critical binding positions within complex structures where key nucleotides and small molecule atoms form hydrogen bonds. This capability demonstrates our model’s ability to capture essential features of genuine binding regions, resulting in accurate predictions of binding events.

Unlike drug development targeting proteins, our understanding of RNA structures is limited, whether through experimental or computational methods, obtaining high-resolution tertiary structure of RNAs is challenging. For different drug design scenarios, we might need to employ various computational virtual screening methods to accelerate drug discovery. On the one hand, for structured molecules like most proteins and certain noncoding RNAs (e.g., Riboswitch and Ribozyme), SBVS methods is suitable. Recently, numerous RNA structure prediction models have been proposed[61, 62], we anticipate that computational predictions of RNA structures will become increasingly accurate, thereby advancing research in structure-based RNA-targeting drug discovery. On the other hand, for RNAs without stable tertiary structures (e.g., many mRNAs and lncRNAs), there remains a lack of prediction methods for RNA-ligand interaction scoring. Disney et al. introduced sequence-based concept by a lead identification screening method which was applied to all human microRNA hairpin precursors[20], in alignment with this advancement, we propose RNAsmol model which provides reliable RNA-small molecule interaction binding prediction without requiring structural input. Our model represents a substantial step forward in leveraging sequence-based approaches to advance the understanding and development of therapeutics targeting RNA interactions. We envision that this deep learning model can serve as a predictive tool to accelerate the development of therapeutic drugs targeting RNA. Additionally, many machine learning-based scoring models for protein-ligand binding suffer from a bias where they memorize molecules rather than learn interactions[51, 58, 63]. This is often due to the advanced deep learning modules that extensively extract features from the molecules themselves but overlook the interaction networks. We have observed a similar issue in RNA-ligand interaction scoring models (**Figure S1**), where predictions under network perturbation yield suboptimal results across various models. Moving forward, our focus will be on addressing this issue to further improve and refine these models or uncover underlying reasons, aiming to enhance the methodological robustness of this research.

## Methods

### Data collection and preprocessing

We initially collected RNA-ligand complex structures from the PDB database, encompassing both RNA-only and RNA-protein (RNP) complexes, to train RNAsmol. Meanwhile, we obtained RNA-small molecule interaction matrices from the ROBIN database which is the largest fully public dataset derived from small molecule microarray (SMM) screening experiments. From the PDB, we gathered experimental RNP-ligand and RNA-ligand complexes with interactions within 4 angstroms, retaining RNP-ligand structures only if RNA atoms constituted more than fifty percent of the total. After applying these filters, 1,229 RNP-ligand and 836 RNA-ligand structures are kept for further screening. Ligands with “non-drug-like” properties were removed adhering to the criteria specified in the referenced paper [64], and we retained only ligands with a mass between 200 and 700 Da. We further filtered RNP-ligand structures to ensure the RNA fraction of the binding sites exceeded 50%. Ultimately, we retained 383 RNP-ligand and 225 RNA-ligand complex structures for extracting chain sequences and small molecule SMILES. All structures were annotated according to their RNA type by text-mining the corresponding PDB file. To validate the effectiveness of RNAsmol using experimental screening data, as shown in **Figure S18**, we compiled SMM screening data from the ROBIN dataset. In this context, we used the hit and non-hit molecules for each RNA target in the screening hit matrix as positive and negative interactions. Basic statistics of these two datasets are shown in **Figure S13, Figure S14, Figure S15**.

### Three perturbations on RNA-small molecule interaction network

For curated raw RNA-small molecule interaction network, there exist three types of relationships between two interacting entities: binding, non-binding, and unknown. Our current knowledge only allows us to determine the molecules that interact with each other, but it fails to establish clear boundaries between non-binding and unknown relationships. To enhance our understanding of this interaction network, we employed various data perturbation strategies to generate non-binding samples from unknown interacting space, as illustrated in **Figure S16**. Firstly, we generated non-binding cases by perturbation on RNA targets through random dinucleotide shuffling and pairing the shuffled sequence with the original small molecules. We denote this kind of perturbation as *ρ*_*r*_:

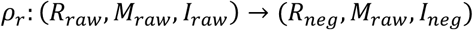

Secondly, we utilized small molecules from different compound libraries (e.g., experimentally validated protein-binder compound libraries, structurally diverse compound libraries, organic small molecule databases) as negative examples for small molecules, where these molecules interact with the original RNA targets to form negative interaction pairs. We denote this kind of perturbation as *ρ*_*m*_:

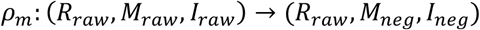

Thirdly, we established edges between each RNA and each small molecule, removed the known edges, and randomly sampled from the remaining edge set to obtain negative example sets. We denote this kind of perturbation as *ρ*_*m*_:

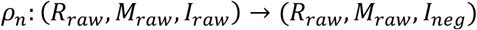

Where *R*_*raw*_ represent raw RNA target set, *M*_*raw*_ represent raw molecule set, *I*_*raw*_ represent the raw RNA-small molecule interaction set, and *R*_*neg*_, *M*_*neg*_ represent the negative RNA targets and negative molecules respectively. We obtain *I*_*neg*_, i.e., the final negative samples for the classification from all of the three perturbations. These three methods perturbed data for both types of interacting entities and the interaction network, aiming to infer binding signals and patterns on the sparse network of RNA-small molecule interactions using diverse data perturbation spaces.

### Three data augmentation strategies on RNA-small molecule interaction network

To address the scarcity of known RNA-ligand binding data, we first augment the RNA by using comparative genomics methods to identify natural binding RNA targets that interact with small molecules and have conserved structures. For RNA sequences with experimental interaction data, we perform large-scale searches across recent metagenomic datasets, clustering homologous sequences based on similarity using the Infernal[65] tool. We hypothesize that although these augmented RNA sequences may differ from those in the PDB database at the sequence level, they can still bind small molecules. Next, we augment the chemical space of small molecules binding to RNA targets, assuming that small molecules with similar chemical properties can also bind to RNA targets. We use computational chemistry methods to map small molecules into continuous numerical molecular fingerprints representing their chemical structure and employ Tanimoto fingerprint similarity metrics for comparison between RNA-binders and drug-like molecules. Based on the assumption that similar RNA targets tend to bind to the same small molecule ligands, and similar small molecules tend to bind to the same RNA targets, we further expand the RNA-ligand binding data using these augmented interaction subsets. We note that there are no augmented interactions between augmented RNAs and augmented small molecules, i.e., edges are only added when one of the interaction partners is a true entity in the raw network. See **Supplementary Methods** and **Figure S16, Figure S17** for details. We only augmented the training dataset to boost model performance, while the validation and test data remained unaugmented. We named these models as RNAsmol_noaug, RNAsmol_rnaaug, RNAsmol_molaug, RNAsmol_bothaug, respectively.

### The RNAsmol model architecture

RNAsmol is RNA-small molecule interaction prediction model with network perturbation and data augmentation. As shown in **Figure 1b**, RNAsmol has four modules: RNA target encoder (Multi-view CNN), small molecule encoder (Graph diffusion convolution), feature fusion module (Bilinear attention block) and classification module (MLP).

### Module1: RNA target encoder (Multi-view CNN)

For the augmented RNA target sequences and their interacting small molecule ligands after redundancy removal, molecular representation and feature extraction are performed separately. For RNA, we retained the first 500 nucleotides of each RNA target sequence and utilized a string representation to depict the sequences and predicted base pairing information from the RNAfold software, i.e., {A, U, C, G, A, a, u, c, g}. Uppercase letters represent paired bases, while lowercase letters indicate unpaired bases. After structural prediction and information normalization of RNA targets, we employ multi-view convolutional neural networks for RNA target local feature extraction. The multi-view convolutional neural network (CNN) architecture is specifically designed to capture diverse local patterns within RNA sequences through multiple convolutional layers with different kernel sizes. This network consists of several primary components: (1) Embedding Layer: The RNA sequence is first embedded into a dense vector representation and transformed into a continuous vector space, which is then suitable for convolutional operations. (2) Convolutional and ReLU Layers: The core of the multi-view CNN comprises several Conv1dReLU blocks. Each block performs a one-dimensional convolution followed by a ReLU activation function.

The convolutional layers have varying kernel sizes (e.g., 3, 5, and 7) to capture different patterns and motifs within the RNA sequences. Formally, given an input sequence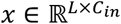, the convolutional operation is defined as:

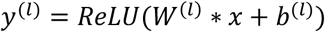

Where *y*^(*l*)^ is the output of the *l*-th convolutional layer, *W*^(*l*)^ and *b*^(*l*)^ are the weights and bias, and * denotes the convolution operation. (3) Stacked CNN Blocks: Multiple StackCNN blocks are used, each containing a stack of convolutional layers with adaptive max pooling. Each block captures features at different levels of abstraction. The stacking of convolutional layers allows the network to learn complex representations from the RNA sequences. (4) Adaptive Max Pooling: After the convolutional operations, adaptive max pooling is applied to reduce the dimensionality of the feature maps, focusing on the most informative features. (5) Feature Aggregation: The outputs from each StackCNN block are concatenated to form a comprehensive feature vector. This aggregated feature vector incorporates diverse local features captured by the different convolutional layers. (6) Fully Connected and Dropout Layers: The concatenated features are passed through a fully connected layer to further integrate the information, followed by a dropout layer to prevent overfitting. This process generates the final feature representation for the RNA target.

### Module2: small molecule encoder (Graph diffusion convolution)

To comprehensively elucidate the binding preferences of small molecules with RNA targets, we adopt atom-level graph representation to encode local features of small molecule ligands. As depicted in **Figure S19**, we initiate by structuring small molecule ligands as graphs, where atoms serve as nodes and chemical bonds as edges. Subsequently, we extract structural and physicochemical features using graph diffusion convolutional neural networks. Traditional graph learning models, often employing Message Passing (MP) methods, typically consider only first-order node neighbors, limiting their ability to abstractly characterize overall graph properties. In contrast, our approach employs a nonlinear information diffusion function to extract features from each point within the molecular graph. This method effectively preserves both high-order local and global graph properties, enhancing feature extraction for RNA binding predictions. Specifically, starting from a fixed atomic node in the small molecule, we conduct graph diffusion based on the transition probability matrix. Upon halting the diffusion process, we define edge weights using the probability distribution from the origin node to other nodes. The graph diffusion process is defined as:

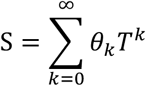

Here, *T* denotes the transition probability matrix, where *T* = *AD*^−1^. *A* represents the adjacency matrix of the molecular graph, and *D* is the degree matrix, with *d*_*ii*_ = ∑_*j*_ *a*_*ij*_. *θ*_*k*_ represents the diffusion coefficient, which commonly includes Personalized PageRank (PPR) diffusion and Heat Kernel (HK) diffusion:

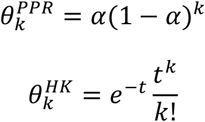

Diffusion convolution, a preprocessing step based on graph diffusion, characterizes the flow of information across the graph structure via random walk processes. This project introduces the novel application of graph diffusion convolutional methods to fully extract feature representations from each atom in the molecular graph. Our goal is to capture comprehensive structural features encoded within molecular graphs and enhance discrimination between molecular graphs of similar small molecules.

### Module3: feature fusion module (Bilinear attention block)

It has been reported that RNA targets and small molecule ligands exhibit diverse binding modes, characterized by specific physicochemical properties and spatial distances. RNA-small molecule binding demonstrates selective specificity, involving various non-covalent interactions like hydrogen bonds and pi-pi stacking, contingent upon interaction strength and physicochemical properties. Traditional models often overlook effective feature fusion, relying solely on simple feature concatenation across layers. In contrast, RNAsmol integrates features from RNA targets and small molecule ligands using a bilinear attention module used in Visual Question Answering (VQA) domain (**Figure S19**). The bilinear attention module contains the following components: (1) Feature transformation: The input RNA features and small molecule features are transformed into the same higher-dimensional space using a fully connected layer respectively. (2) Attention computation: The transformed features are computed to get attention maps either using single-view attention which conducts tensor contraction operation to get attention scores or using multi-view attention which involves creating higher-dimensional tensors and utilize linear transformation to get attention scores. (3) Softmax activation (4) Pooling and Fusion (5) Output. Leveraging attention maps and pooling strategies facilitates the extraction and fusion of relevant information from both modalities and enhance predictive performance and generalization across diverse datasets. As shown in **Figure S4**, ablation studies on bilinear attention network (BAN) module underscore the pivotal role of this fusion module in effectively classifying RNA-small molecule interactions.

### Module4: classification module (MLP)

We used the Multi-Layer Perceptron (MLP) module consisting of three to five fully connected (dense) layers interspersed with Rectified Linear Unit (ReLU) activation functions as the classification module to transform fused feature embedding encoded by the bilinear transformer module into the probability for each label. Generally, each layer *L*_*i*_ in the network applies a linear transformation to its input, followed by a ReLU non-linearity. The linear transformation for a given layer *L*_*i*_ can be represented as:

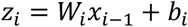

Where *W*_*i*_ and *b*_*i*_ are the weight matrix and bias vector for layer *L*_*i*_ respectively, and *x*_*i*−1_is the output of the previous layer. The ReLU activation function is applied element-wise to the linear transformation output:

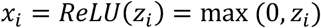

Besides, we employ the Cross-Entropy Loss function (‘nn.CrossEntropyLoss’) provided by PyTorch and ensure the output layer has two neurons corresponding to the two classes. The cross-entropy (CE) loss function is defined as:

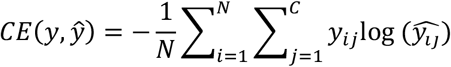

Where *y*_*ij*_ is the true label, 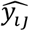 is the predicted probability for class *j* for sample *i, N* is the number of samples, and C is the number of classes (which is 2 in the case of binary classification in RNAsmol). We utilize Adam optimizer configured with the given learning rate and an *L*2 regularization term (weight decay). The weight decay term helps to prevent overfitting by penalizing large weights, thereby improving the generalization capability of the model. The loss curves of model training are displayed in **Figure S20**.

### 10-fold cross-validation (CV) evaluation

To evaluate the classification performance, we performed 10-fold cross-validation on RNAsmol and other related models, comparing them across multiple metrics including ROCAUC, PRAUC, accuracy (ACC), sensitivity (SEN), specificity (SPE) and F1 score. Since there is a lack of binary classification prediction models designed for RNA-small molecule interactions, we modified the molecular encoding part of recently published models for predicting protein-ligand binding interactions to accommodate RNA molecules. These adapted models, which we named GraphDTA_RNA, MGraphDTA_RNA, IIFDTI_RNA and DrugBAN_RNA, are detailed as follows. The GraphDTA model uses a graph neural network to learn small molecule SMILES and a convolutional neural network to learn protein sequences, followed by a fully connected neural network to predict binding probabilities after the simple concatenation of the extracted features. We revised the initial embedding of protein sequences to fit RNA sequences and used the default settings of this model for evaluation. The MGraphDTA model uses a multi-view graph neural network (MGNN) to learn small molecule SMILES and a multi-view convolutional neural network to learn protein sequences, predicting binding probabilities with a fully connected neural network after feature concatenation. We adjusted the initial embedding of protein languages to fit RNA sequences and used the default settings of this model for evaluation. For the IIFDTI model, we modified the embedding module as follows: (1) We replaced the protein text corpus with Rfam[66] and trained a skip-gram model from gensim Word2Vec on it to obtain k-mer embeddings from RNA sequences. (2) We applied the trained rna2vec vector to the RNA target. We then used the default settings in the IIFDTI model for evaluation. For the evaluation of DrugBAN on the RNA-ligand classification task, we used the default parameters provided in DrugBAN.yaml and employed the random split method.

### Cold evaluation

To evaluate the classification performance on unseen datasets, we conducted cold evaluation on RNAsmol and other models mentioned in the previous section. This involved ensuring that the test set included RNA targets, small molecule ligands, and both interacting molecules that had not appeared in the training set. In the cold evaluation on RNA, there is no overlap between RNA targets in the training and testing sets (R_train and R_test in **Figure S18**), though small molecule ligands may overlap (M_train and M_test). This approach trains a model particularly suited for predicting small molecule ligands for new RNA targets of interest. In the cold evaluation on small molecule, there is no overlap between small molecules in the training and testing sets, while RNA targets may overlap. This setting trains a model suitable for predicting appropriate RNA targets for newly discovered or unvalidated small molecule ligands that bind to RNA. The cold evaluation on pair ensures no overlap between both RNA targets and small molecules in the training and testing sets, which is the strictest setting and usually results the least accurate predictions. It is worth noting that, similar to random splitting used in cross-validation evaluations, there is no overlap between interactions in the training and testing sets (I_train and I_test in **Figure S18**). However, specific requirements are made in cold evaluations for selecting the two interacting molecules. With these methods, we can apply the trained RNA-small molecule binding prediction model to practical prediction scenarios, aiming to discover potential small molecule drug sets for specific RNA targets or predict appropriate RNA targets for small molecule drugs.

### Decoy evaluation

To reveal the potential of virtual screening in RNA-targeting drug discovery, we did 10-fold decoy evaluation on RNAsmol and other models including RNAmigos, RNAmigos2 and rDock. First, we generated a decoy set for each small molecule in the test sets of PDB dataset using DecoyFinder software on ZINC bioactive library. This software selects molecules with similar physicochemical properties (including molecular weight (MW), partition coefficient (logP), hydrogen bonds donors (HBD), hydrogen bond acceptors (HBA) and number of rotatable bonds (RB)) but not too similar molecular structures from the given library for each query molecule. Then, we used the decoy sets for model evaluation and comparison. For RNAsmol, we ranked the predicted binding score of the true RNA-binder within the predicted scores of all molecules in its decoy set, a higher rank indicates a better decoy rank score. For RNAmigos, we generated a predicted fingerprint using a trained model and ranked all compounds in the decoy set according to their distance from the predicted fingerprint. We calculated fingerprint similarity using four distance metrics: Euclidean distance, cosine distance, Chebyshev distance and correlation distance, with differences shown in **Figure S8**. For RNAmigos2, we used four modes of this model including dock mode, native mode, fp mode and mixed mode for the evaluation, directly ranking the predicted score within the decoy as the decoy rank score. For rDock, we used the docking scores from the default outputs for the ranking and evaluation. Moreover, we performed a target shuffle in the decoy evaluation to disclose the RNA target specificity and robustness of the models. Instead of generating a brand-new decoy set, we shuffled the correspondence between RNA targets and the decoy sets through random sampling and reran the four models.

To generalize the decoy evaluation to unseen data, we applied trained RNAsmol_PDB and RNAsmol_ROBIN model, incorporating *ρ*_*m*_ perturbation, to identified RNA-targeting drugs such as ribocil and risdiplam. And we used RNAmigos, RNAmigos2 and RSAPred_riboswitch models for decoy evaluation and comparison. First, we generated decoy sets for each small molecule and used the trained models to calculate decoy rank scores for comparison. We converted the compounds to SMILES format using computational tools such as rdkit and mathpix OCR. The 3D structures of ribocil-targeted RNA, risdiplam-targeted RNA, MALAT1 RNA were curated from the PDB database. We used two chains in the FMN riboswitch structure (PDB ID: 5KX9), the 5’-end of U1 snRNA and the 5’-splice sites of the SMN2 exon7 structure (PDB ID: 6HMO), as well as the MALAT1 triple helix structure (PDB ID: 4PLX) as target sequences for the prediction in RNAsmol. For RNAmigos and RNAmigos2, we extracted the pockets from the available structures using the molecule-binding positions in the complexes or positions mentioned in the literature. Finally, we reported the best rank score of the two chains in our results.

### Case study validation

To better interpret the prediction of the RNAsmol model, we use the gradient-weighted class activation mapping (Grad-CAM) algorithm, which employs gradients backpropagated from the prediction layer to the activations of interest. We focus on the last convolutional layers of multi-view convolutional neural networks in the RNA encoder and the graph diffusion neural networks in the small molecule encoder to illustrate the weights on individual nucleic acids and atoms. These weights are displayed on the RNA secondary structure plot using forna[67] visualization tool and the small molecular graph drawn by rdkit (https://www.rdkit.org) (**Figure 6** and **Figure S9**). Besides, we visualize the interaction profile of PreQ1, ZTP, SAH, and c-di-GMP-II riboswitch in 2D and 3D complex structures using Ligplot+[68] and PyMOL[69] (**Figure 6, Figure S10, Figure S11**). The position of hydrogen bonds is annotated in the structure and profile using the functions in these tools, with default hydrogen-bond calculation parameters in Ligplot+ set to a maximum hydrogen-acceptor distance of 6 and a minimum acceptor-donor distance of 6.

## Declarations

### Data availability

All datasets used in this study are publicly available for academic use.

## Author contributions

H.M., and Z.J.L. conceived and designed the project. H.M., Y.J., K.L. completed the preprocessing of the data. H.M., Y.B. developed the framework of the model and performed the experiments. H.M., L.G. performed the evaluation of the model and analyses. H.M.wrote the manuscript. Z.J.L., Y.J., X.L., P.B. revised the manuscript.

